# Activation of the Extracytoplasmic Function σ factor σ^P^ by β-lactams in *Bacillus thuringiensis* requires the site-2 protease RasP

**DOI:** 10.1101/707133

**Authors:** Theresa D. Ho, Kelsie M. Nauta, Ute Müh, Craig D. Ellermeier

## Abstract

Bacteria can utilize alternative σ factors to regulate sets of genes in response to changes in the environment. The largest and most diverse group of alternative σ factors are the Extracytoplasmic Function (ECF) σ factors. σ^P^ is an ECF σ factor found in *Bacillus anthracis*, *B. cereus*, and *B. thuringiensis*. Previous work showed σ^P^ is induced by ampicillin, a β-lactam antibiotic, and required for resistance to ampicillin. However, it was not known how activation of σ^P^ is controlled or what other antibiotics may activate σ^P^. Here we report that activation of σ^P^ is specific to a subset of β-lactams and σ^P^ is required for resistance to these β-lactams. We demonstrate that activation of σ^P^ is controlled by the proteolytic destruction of the anti-σ factor, RsiP, and that degradation of RsiP requires multiple proteases. Upon exposure to β-lactams, the extracellular domain of RsiP is cleaved by an unknown protease, which we predict cleaves at site-1. Following cleavage by the unknown protease, the N-terminus of RsiP is further degraded by the site-2 intramembrane protease, RasP. Our data indicate that RasP cleavage of RsiP is not the rate-limiting step in σ^P^ activation. This proteolytic cascade leads to activation of σ^P^ which induces resistance to β-lactams likely via increased expression of β-lactamases.

**Importance:** The discovery of antibiotics to treat bacterial infections has had a dramatic and positive impact on human health. However, shortly after the introduction of a new antibiotic bacteria often develop resistance. The bacterial cell envelope is essential for cell viability and is the target of many of the most commonly used antibiotics including β-lactam antibiotics. Resistance to β-lactams is often dependent upon β-lactamases. In *B. cereus*, *B. thuringiensis* and some *B. anthracis* strains the expression of some β-lactamases is inducible. This inducible β-lactamase expression is controlled by activation of an alternative σ factor called σ^P^. Here we show that β-lactam antibiotics induce σ^P^ activation by degradation of the anti-σ factor RsiP.

## Introduction

The bacterial cell envelope is essential for cell viability and is the target of many of the most commonly used antibiotics including β-lactams like penicillins, penems, and cephalosporins. These are broad-spectrum antibiotics that target peptidoglycan biosynthesis by inhibiting the transpeptidase activity of penicillin-binding proteins. This results in decreased and/or altered crosslinking of peptidoglycan which leads to cell envelope damage and subsequent cell lysis and death (1, 2).

Members of the *Bacillus cereus* group, including *Bacillus thuringiensis* and *Bacillus cereus* and some strains of *Bacillus anthracis*, are highly resistant to β-lactam antibiotics (3–6). This resistance is due in part to expression of at least two β-lactamases (3, 5). The expression of these β-lactamases is induced by ampicillin and is dependent upon the alternative σ factor, σ^P^. σ^P^ belongs to the Extracytoplasmic Function (ECF) family of alternative σ factors (5).

Bacteria often utilize alternative σ factors to regulate subsets of genes required for survival in specific environmental conditions or stress responses. ECF σ factors are the largest and most diverse group of alternative σ factors and represent the “third pillar” of bacterial signal transduction (7, 8). ECF σ factors belong to the σ^70^ family but unlike the “housekeeping” σ factor, σ^70^, ECF σ factors contain only region 2 and region 4.2 of σ^70^ which recognize and bind to the −10 and −35 regions of promoter sequences, respectively (8, 9). In addition, unlike σ^70^, ECF σ factors are generally held inactive by anti-σ factors until bacteria encounter an inducing signal (10, 11). Upon induction, ECF σ factors are released from their cognate anti-σ factors to promote transcription of specific stress-response genes.

The ECF σ factors have been subdivided into more than 40 distinct groups with ECF01 being the best studied [Reviewed in (7, 11, 12)]. σ^P^ belongs to the ECF01 family which includes members like σ^E^ and σ^W^ from *E. coli* and *Bacillus subtilis*, respectively. The activities of the ECF01 family are inhibited by their cognate transmembrane anti-σ factors (8, 13). To activate ECF01 σ factors, the anti-σ factors must be destroyed via a proteolytic cascade (14, 15). For example, the *E. coli* anti-σ factor, RseA, is degraded in response to outer membrane stress leading to σ^E^ activation (16, 17). DegS, a serine protease, cleaves the anti-σ factor RseA at site-1 (14, 18, 19). After site-1 cleavage, the conserved site-2 protease, RseP, cleaves RseA within the membrane leading to increased σ^E^ activity (14, 20, 21). Similarly, the σ^W^ anti-σ factor, RsiW, from *B. subtilis* is proteolytically degraded by site-1 and site-2 proteases. In the case of RsiW, the site-1 protease is PrsW, a metalloprotease unrelated to DegS. PrsW cleaves RsiW in response to antimicrobial peptides, vancomycin and pH change (22–24). RsiW is further processed by the conserved site-2 protease RasP, a homolog of RseP (15).

The closely related ECF30 family member, σ^V^ from *B. subtilis*, is activated by lysozyme (25–29). Activation of σ^V^ differs from σ^E^ and σ^W^ activation in that σ^V^ is not controlled by a dedicated site-1 protease but instead utilizes signal peptidases (30, 31). Signal peptidases are essential proteases which are required to cleave substrates secreted from the general secretion or twin arginine secretion systems (32–34). The anti-σ factor, RsiV binds to lysozyme which allows signal peptidase to cleave RsiV at site-1 (30, 31). This allows the site-2 protease, RasP, to cleave RsiV, leading to σ^V^ activation (35).

Previous studies found σ^P^ is induced by ampicillin and its activity is required for resistance to ampicillin (5). The activity of σ^P^ is inhibited by the transmembrane anti-σ factor, RsiP (5, 6). However, whether σ^P^ is activated specifically by ampicillin or more generally by cell wall stress is not known. In *B. subtilis*, activation of σ^V^ is specific to lysozyme (26, 27), while activation of σ^W^, σ^X^ and σ^M^ is in response to more general cell envelope stress (9, 36, 37). Here, we show σ^P^ is activated by a specific subset of β-lactams and this activation occurs via regulated intramembrane proteolysis of the anti-σ factor, RsiP.

## Results

### A subset of β-lactams induces σ^P^ activation

Previously, Koehler and colleagues demonstrated that ampicillin induces expression of the β-lactamase encoded by *bla1* (*hd73_3490*) in a σ^P^-dependent manner in *B. thuringiensis* and *B. cereus* (5). Activation of some ECF σ factors is highly specific to an inducing signal, while others are activated by more general cell envelope stress. Thus, we sought to determine the specificity of σ^P^ activation using *B. thuringiensis* as a model system.

Like many ECF σ factor systems, σ^P^ is required for its own transcription (5). To monitor σ^P^ activation, we fused the σ^P^ promoter (P_*sigP*_) to the *lacZ* reporter gene and integrated this construct into the genome of *B. thuringiensis* (THE2549 *thrC*::P_*sigP*_-*lacZ*). We tested several classes of β-lactams and cell wall-targeting antibiotics for their ability to induce expression of P_*sigP*_-*lacZ*. We observed wide zones of P_*sigP*_-*lacZ* induction around cefoxitin and cefmetazole (Fig. 1). We detected fainter zones of induction in the areas around cephalothin and cephalexin (Fig. 1). Very faint zones of induction were present in the cells around ampicillin and methicillin (Fig.1). Interestingly, we did not observe this induction surrounding the β-lactams, cefoperazone and piperacillin or antibiotics that target other steps in cell wall biosynthesis including, ramoplanin, phosphomycin, nisin, bacitracin, and vancomycin (Fig. 1). We also tested compounds that do not target peptidoglycan biosynthesis including kanamycin, polymyxin B and erythromycin and saw no induction of P_*sigP*_-*lacZ* (Fig.1).

**Figure 1.**
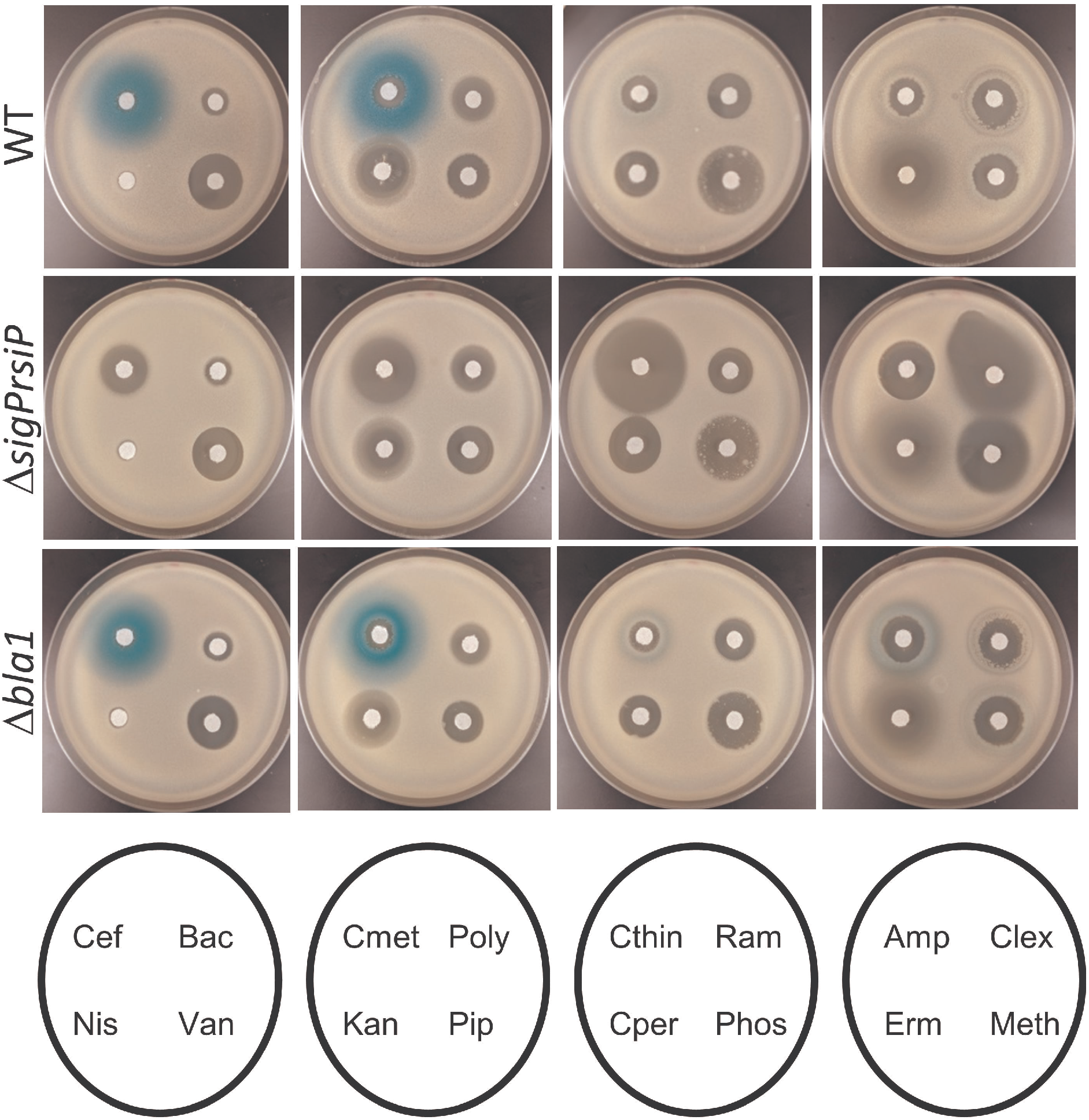
Expression of *sigP* is specifically induced by β-lactams. All the strains contained P_*sigP*_-*lacZ* in either wild type (THE2549), Δ*sigPrsiP* (EBT232) or Δ*blaI* (EBT215) background. Mid-log cells were washed and diluted 1:100 in molten LB agar containing X-gal (100 μg/ml) and poured into empty 100 mm Petri dishes. Filter disks containing Cef (1 μl of 5 mg/ml cefoxitin), Bac (1 μl of 50 mg/ml bacitracin), Nis (3 μl of 100 mg/ml nisin), Vanc (1 μl of 10 mg/ml vancomycin), Cmet (1 μl of 5 mg/ml cefmetazole), Poly (1 μl of 50 mg/ml polymyxin B), Kan (1 μl of 10 mg/ml kanamycin), Pip (1 μl of 5 mg/ml piperacillin), Cthin (1 μl of 50 mg/ml cephalothin), Ramo (1 μl of 25 mg/ml ramoplanin), Cper (1 μl of 50 mg/ml cefperazone), Phos (1 μl of 100 mg/ml phosphomycin), Amp (2 μl of 200 mg/ml ampicillin), Clex (1 μl of 50 mg/ml cefalexin), Erm (1 μl of 5 mg/ml erythromycin), and Meth (2 μl of 100 mg/ml methicillin) were then placed on the top agar and incubated for 16 hours at 30°C.

To quantify the levels of β-lactam induction, we tested eight β-lactams for their ability to activate the P_*sigP*_-*lacZ* fusions using a β-galactosidase assay. Mid-log cells were incubated in the presence of various concentrations of ampicillin, cefoxitin, cefmetazole, cephalothin, methicillin, cephalexin, cefoperazone, and cefsulodin for 1 hour at 37°C. We observed dose-dependent induction with a subset of these β-lactams (Fig. 2A-B). Interestingly, ampicillin, methicillin and cephalexin showed low levels of P_*sigP*_-*lacZ* induction when spotted onto a lawn of cells (Fig. 1) but strongly induced P_*sigP*_-*lacZ* in liquid assays (Fig. 2A-B), a point we will return to later. In contrast, neither cefoperazone nor cefsulodin were able to induce on the plates or in liquid (Fig. 1 and 2B). This confirms our observation that a sub-set of β-lactams induce σ^P^ activation.

**Figure 2.**
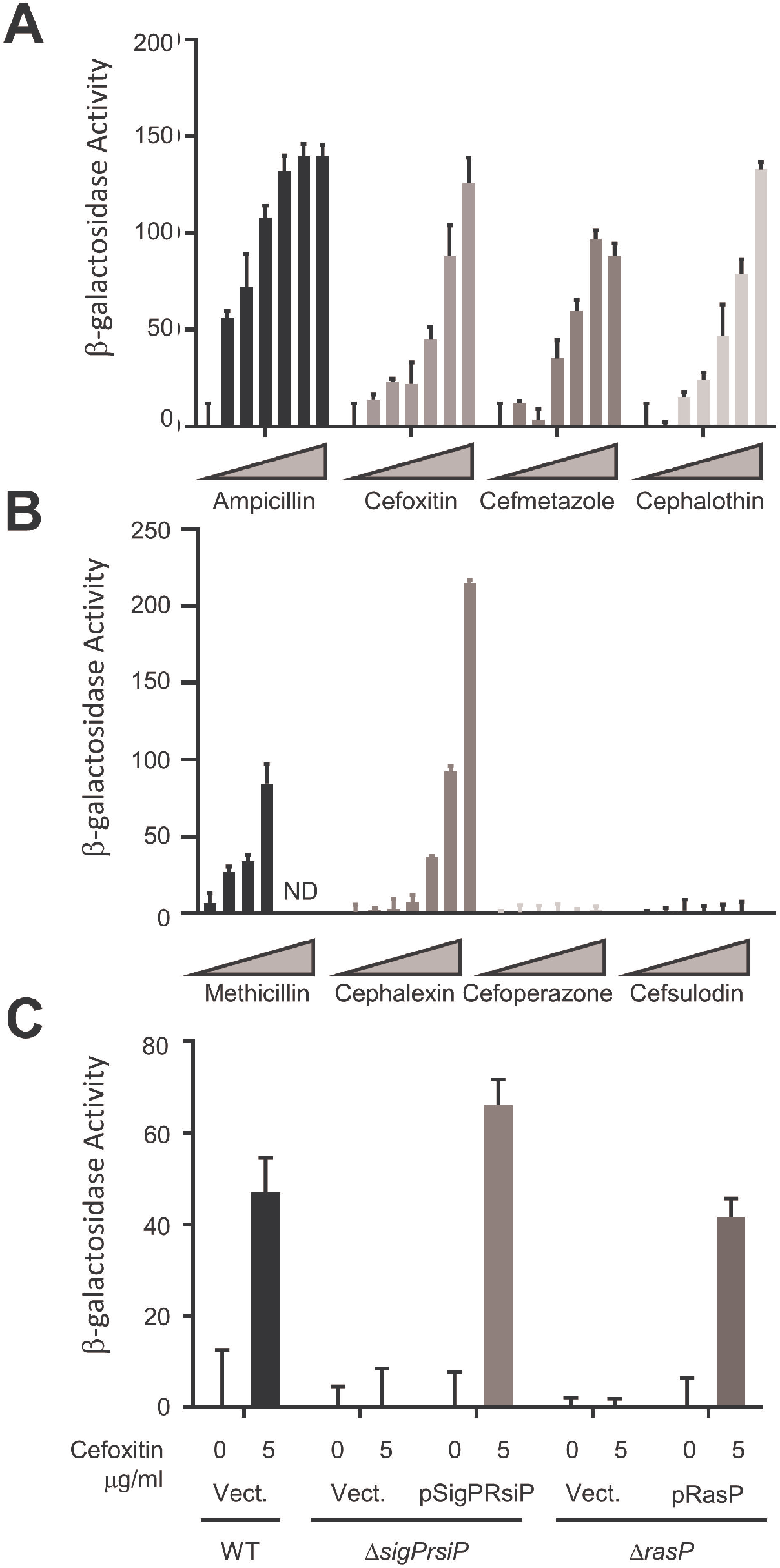
Expression of P_*sigP*_-*lacZ* is dose-dependent and dependent upon σ^P^ and RasP. **A.** *B. thuringiensis* with a transcriptional fusion P_*sigP*_-*lacZ* (THE2549) was grown overnight at 30°C and subcultured in LB and grown to OD_600_ ~0.8 before being incubated with varying concentrations of β-lactams (0, 0.0625, 0.125, 0.25 0.5, 1, and 2 μg/ml) for 1 hour. Cells were collected and resuspended in Z-buffer. **B.** *B. thuringiensis* with a transcriptional fusion P_*sigP*_-*lacZ* (THE2549) was grown overnight at 30°C and subcultured in LB and grown to OD_600_ ~0.8 before being incubated with varying concentrations of β-lactams (0, 0.0625, 0.125, 0.25 0.5, 1, and 2 μg/ml) for 1 hour. Cells were collected and resuspended in Z-buffer. **C.** All strains contain P_*sigP*_-*lacZ* and the genotype and plasmid noted: wild type/Vect (EBT169); *sigP*/Vect (EBT251); Δ*sigPrsiP*/pSigPRsiP (EBT238); Δ*rasP*/Vect (EBT175); *rasP*/pRasP (EBT176). Strains were grown to mid-log then treated with cefoxitin 5 μg/ml (5) or untreated (0) and incubated for 1 hour. β-Galactosidase activity was calculated as described in the material and methods. These experiments were done in triplicate and standard deviation is represented by error bars.

We found that deletion of the *sigPrsiP* genes blocked expression of P_*sigP*_-*lacZ* in the presence of β-lactams (Fig. 1 and Fig. 2C) demonstrating σ^P^ is required for induction of P_*sigP*_-*lacZ* in response to β-lactams. When we introduced a low copy plasmid containing P_*sigP*_-*sigP*^+^*rsiP*^+^ into the Δ*sigPrsiP* mutant (Δ*sigPrsiP*/pSigPRsiP), we restored the induction of P_*sigP*_-*lacZ* in response to cefoxitin (Fig. 2C). Taken together, these data suggest a subset of β-lactam antibiotics activate σ^P^.

### σ^P^ and Bla1 are involved in resistance to some β-lactams

To determine the impact of σ^P^ on resistance to β-lactams, we measured the minimal inhibitory concentration (MIC) of wild type and a Δ*sigPrsiP* mutant for several β-lactams. We found the wild type was greater than 100-fold more resistant to ampicillin, methicillin, and cephalothin than was the Δ*sigPrsiP* mutant (Table 1). Wild type was 16 to 50-fold more resistant to cefmetazole, cefoxitin and cephalexin than the mutant (Table 1). There was little or no difference in resistance to piperacillin, cefoperazone and cefsulodin which also failed to activate σ^P^ (Table 1 and Fig. 1). We also demonstrate that complementing the Δ*sigPrsiP* mutant with a plasmid carrying P_*sigP*_-*sigP*^+^-*rsiP*^+^ restored resistance to ampicillin and cefoxitin (Table 2). For reasons that remain unclear, strains containing plasmids including empty vector have slight increases in β-lactam resistance. However, this does not impact the observation that the presence of P_*sigP*_-*sigP*^+^-*rsiP*^+^ restored resistance to ampicillin and cefoxitin.

**Table 1.** *ΔsigPrsiP* is more sensitive to β-lactams than WT.

|  | **WT<sup>1</sup>** | ***ΔsigPrsiP*<sup>1</sup>** | **Fold Difference** |
| --- | --- | --- | --- |
| Ampicillin | 6000 +/- 0 | 1.67 +/- 0.5 | 3592 |
| Cefoxitin | 200 +/- 0 | 20 +/- 0 | 10 |
| Methicillin | 666 +/- 115 | 1 +/- 0 | 666 |
| Piperacillin | 5 +/- 0 | 1.25 +/- 0 | 4 |
| Cephalothin | 88 +/- 25 | 0.25 +/- 0 | 350 |
| Cephalexin | 200 +/- 0 | 4 +/- 0 | 50 |
| Cefmetazole | 44 +/- 13 | 2.8 +/- 1.1 | 16 |
| Cefoperazone | 5 +/- 2 | 4 +/- 0 | 1.25 |
| Cefsulodin | 400 +/- 0 | 400 +/- 0 | 1 |

<sup>1</sup> MIC of antibiotic concentrations reported as μg/ml

**Table 2.**
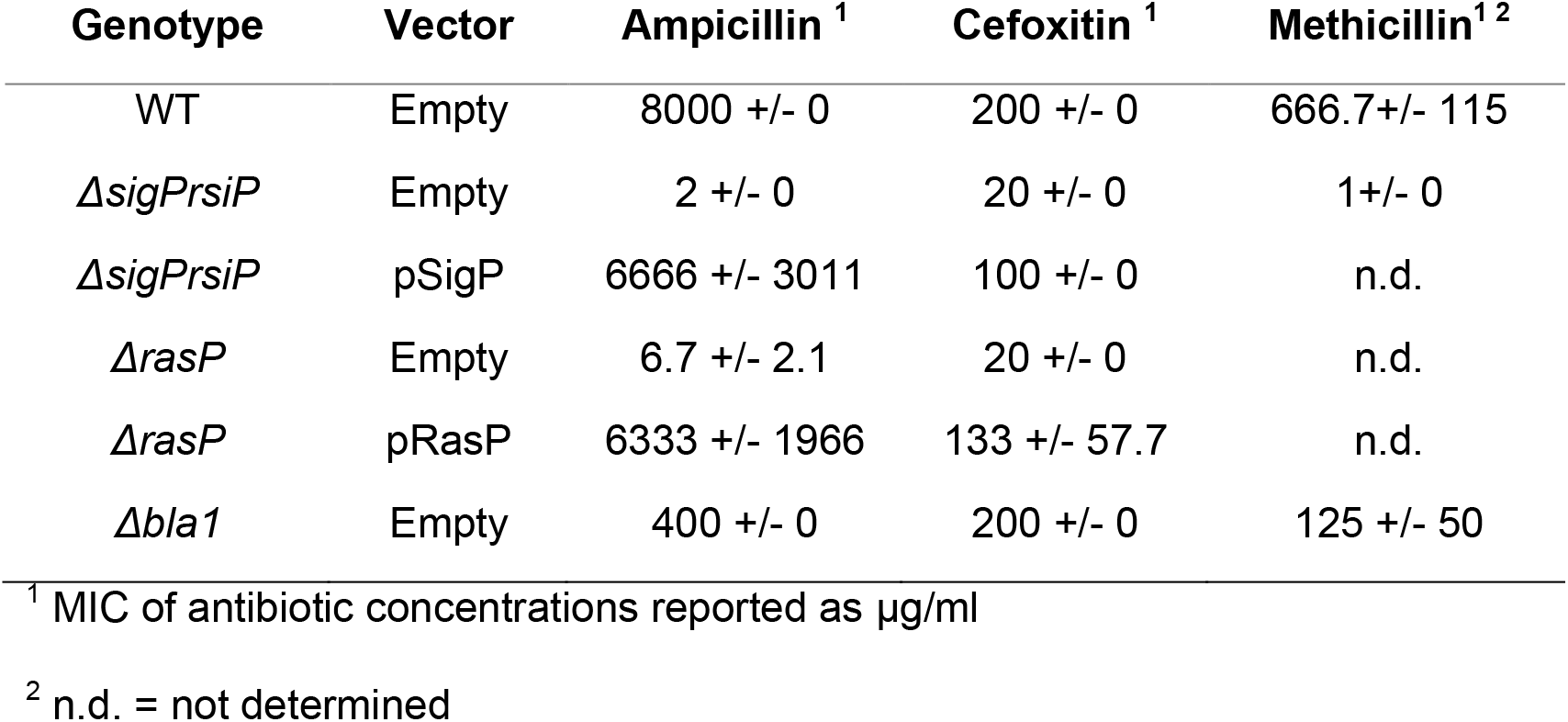
RasP is required for resistance to β-lactams.

Since σ^P^ was shown to control expression of *hd73_3490* (referred to hereafter as *bla1*), which encodes a β-lactamase, we sought to determine if this gene played a role in resistance to β-lactams. We made a deletion of *bla1* and determined the MIC of ampicillin and cefoxitin for this strain. The *bla1* mutant was 8 to 16-fold more sensitive to ampicillin, ~5 fold more sensitive to methicillin but no more sensitive to cefoxitin than wild type (Table 2). This contrasts with the *sigP* mutant which is greater than 1000-fold more sensitive to ampicillin, 600-fold more sensitive to methicillin and ~25-fold more sensitive to cefoxitin than the wild type (Table 2). This suggests that Bla1 plays a more important role in resistance to ampicillin and methicillin than to cefoxitin. Furthermore, our data suggests that while Bla1 contributes to β-lactam resistance, additional σ^P^ regulated genes must also contribute to β-lactam resistance.

When we tested various β-lactams for induction of P_*sigP*_-*lacZ* on X-gal plates, we did not consistently observe a strong zone of induction surrounding ampicillin and methicillin (Fig. 1). We hypothesized this weak induction zone was because the wild type efficiently produced β-lactamases which degraded the inducer (ampicillin and methicillin). Thus, we were unable to observe the increased production of β-galactosidase. To test this hypothesis, we determined the effect of a Δ*bla1* mutant on σ^P^ activation. We found that in the Δ*bla1* mutant, ampicillin and methicillin produced more distinct zones of induction (Fig. 1). However, all other induction zones of the Δ*bla1* mutant were similar to wild type. Thus, in the absence of Bla1 which degrades ampicillin and methicillin, we detected greater induction of P_*sigP*_-*lacZ* expression. Taken together, these observations suggest that the weak ampicillin induction of P_*sigP*_-*lacZ* on plates is in part due to the efficient degradation of the inducer by β-lactamases.

### RsiP is degraded in response to cefoxitin in a dose-dependent manner

The anti-σ factors of other ECF01 family members are degraded which leads to the activation of their cognate σ factors (7, 14, 15). We sought to determine if β-lactams activate σ^P^ by inducing degradation of RsiP. To investigate this, we constructed a strain with an anhydrotetracycline-inducible copy of green fluorescent protein (GFP) fused to the N-terminus of RsiP (GFP-RsiP). The inducible promoter allows us to uncouple expression of RsiP from induction of σ^P^. The GFP-RsiP fusion allows us to follow the fate of the cytoplasmic portion of RsiP. Expression of GFP-RsiP complements an *rsiP* null mutation (Fig. S1) and localizes to the membrane (Fig. S2). We then induced the synthesis of GFP-RsiP in exponential phase cells and monitored its processing before and after treatment with cefoxitin. We chose to utilize cefoxitin for these experiments because cefoxitin induces σ^P^ activation over a wide concentration range and the *ΔsigPrsiP* mutant strain grows at most of these concentrations (Fig. 2A and Table 1). Cell pellets were then lysed by sonication and western blots were performed using anti-RsiP antisera against the extracellular portion of RsiP or anti-GFP antisera which detects GFP fused to the intracellular portion of RsiP.

When cells producing GFP-RsiP were grown in the absence of cefoxitin we detected full length GFP-RsiP at the expected size of ~60 kD using anti-RsiP anti-sera. This band was absent in the empty vector control (Fig. 3A). When cells were incubated with cefoxitin (5 μg/ml) for various times we found that the level of full-length GFP-RsiP decreased over time (Fig. 3A and Fig. S3A). We observed loss of GFP-RsiP by 30 minutes to 1-hour post exposure to cefoxitin (Fig. 3A and Fig. S3A). This suggests GFP-RsiP is likely degraded in the presence of cefoxitin.

**Figure 3.**
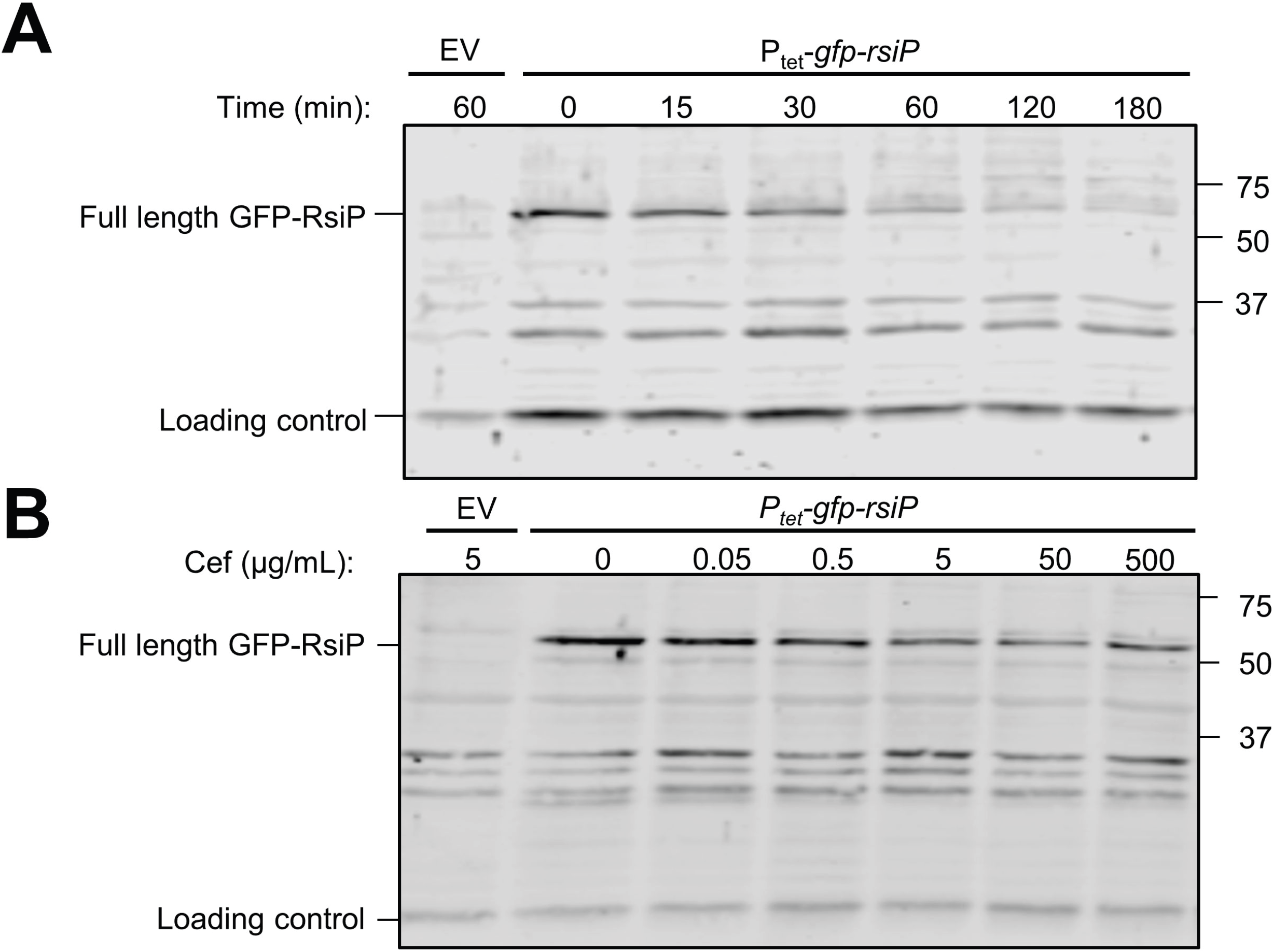
RsiP levels decrease in the presence of cefoxitin. *B. thuringiensis* expressing tetracycline-inducible *gfp-rsiP* (EBT360) or empty vector (EV; EBT169) was subcultured 1:50 into LB supplemented with ATc (50 ng/ml). At mid-log cells were incubated with **A.** 5 μg/ml of cefoxitin for various times (0, 15, 30, 60, 120, or 180 minutes) or **B.** increasing concentrations of cefoxitin (0, 0.05, 0.5, 5, 50, 500 μg/ml) for 1 hour. The immunoblot was probed with antisera against RsiP (α-RsiP^76-275^). Streptavidin IR680LT was used to detect HD73_4231 (PycA homolog) which served as a loading control (69, 70). The color blot showing both anti-RsiP and Streptavidin on a single gel is Fig. S3.

We also tested the effect of cefoxitin concentration on GFP-RsiP levels by incubating cells with a range of cefoxitin concentrations (0-500 μg/mL) for one hour. We found increasing concentrations of cefoxitin resulted in greater decrease of full length GFP-RsiP (Fig. 3B and Fig. S3B). We obtained comparable results when we blotted for the N-terminal domain using anti-GFP antisera (Fig. S4). These data suggest activation of σ^P^ occurs via loss of RsiP in a cefoxitin dose-dependent manner.

### RasP is necessary for σ^P^ activation

Both σ^E^ and σ^W^ are activated by regulated intramembrane proteolysis of their cognate anti-σ factors. Proteolysis of these anti-σ factors requires multiple proteases, including the highly-conserved site-2 protease, RseP and RasP, respectively (14, 15). We hypothesize that activation of σ^P^ requires multiple proteases including the conserved site-2 protease RasP to degrade RsiP. To test this we used BLAST to identify a putative membrane embedded metalloprotease, HD73_4103, which is 76% similar and 60% identical to *B. subtilis* RasP and is hereafter referred to as RasP (Fig. S5) (38–43). To determine if RasP was required for σ^P^ activation, we generated a strain containing a deletion of *rasP* and the P_*sigP*_-*lacZ* reporter. In the absence of RasP, we do not detect increased expression of P_*sigP*_-*lacZ* reporter in response to cefoxitin (Fig. 2C). In MIC experiments, we found that, similar to the Δ*sigPrsiP* mutant, the Δ*rasP* mutant was more sensitive to ampicillin and cefoxitin (Table 2). We found both resistance to β-lactams and induction of P_*sigP*_-*lacZ* could be complemented when a plasmid expressing *rasP*^+^ was introduced into the Δ*rasP* mutant (Fig. 2C and Table 2). These data suggest RasP is required for σ^P^ activation.

### RasP is required for degradation of RsiP

To determine if RasP is required for degradation of RsiP, we expressed the GFP-RsiP fusion in both wild type and a Δ*rasP* mutant. We treated cells with 5 μg/mL cefoxitin for various lengths of time from 0 to 180 minutes (Fig. 4 and Fig. S6). In wild type, we observed loss of full-length RsiP over time (Fig. 4 and Fig. S6). In contrast, we observed loss of full-length GFP-RsiP and the accumulation of a smaller ~35kD band in the Δ*rasP* mutant (Fig. 4 and Fig. S6). This suggests RasP is required for complete degradation of RsiP. Since a truncated product accumulates in the Δ*rasP* mutant, RasP is likely required for site-2 cleavage and an unidentified protease is required for cleavage at site-1.

**Figure 4.**
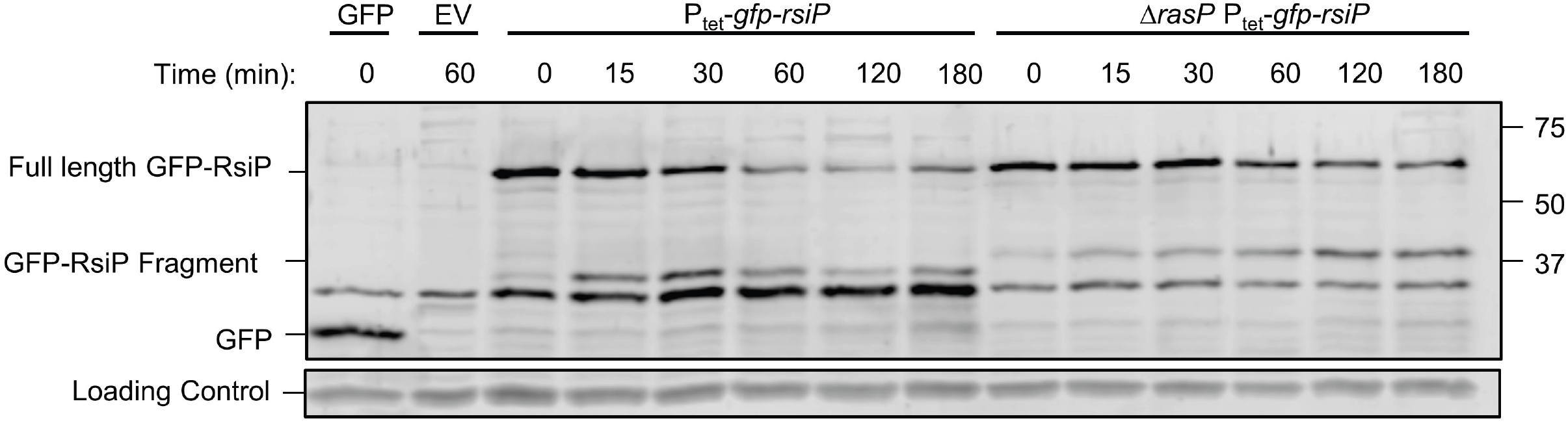
RsiP degradation is dependent upon the site-2 protease RasP. *B. thuringiensis* wild type (EBT360) or *ΔrasP* (EBT366) containing a tetracycline-inducible copy of *gfp-rsiP* were subcultured 1:50 into LB supplemented with ATc (50 ng/mL). At mid-log cultures were incubated for 1 hour untreated (−) or treated (+) with cefoxitin (5 μg/ml) at 37°C. The immunoblot was probed with anti-GFP antisera. EV is wild type with pAH9 (EBT169) and GFP is wild type with pAH13 (UM20). Streptavidin IR680LT was used to detect HD73_4231 (PycA homolog) which served as a loading control (63, 64). The color blot showing both anti-GFP and Streptavidin on a single gel is Fig. S6.

### Mutations in *rsiP* result in constitutive *sigP* expression

To further characterize the σ^P^ signal transduction system, we isolated mutants which resulted in constitutive expression of P_*sigP*_-*lacZ*. We selected for mutants with increased resistance to cefoxitin by plating cultures of the wild type P_*sigP*_-*lacZ* strain (THE2549) on LB cefoxitin 200 μg/ml agar. At this concentration of cefoxitin, wild type *B. thuringiensis* fails to grow. These strains were tested for P_*sigP*_-*lacZ* expression in the absence of cefoxitin by streaking on LB X-gal. We isolated 8 independent mutants with increased resistance to cefoxitin that have constitutive P_*sigP*_-*lacZ* expression. We hypothesized these strains harbored mutations in *rsiP*. We PCR amplified and sequenced the *sigP* and *rsiP* genes from the constitutive mutants. The 8 constitutive mutants contained mutations in different regions of the *rsiP* gene that resulted in C-terminal truncations of RsiP (Fig. S7). We selected four *rsiP* mutants for further study. We found that each mutant strain showed increased P_*sigP*_-*lacZ* expression even in the absence of β-lactams (Fig. 5). When a wild type copy of *rsiP* (pSigPRsiP) was introduced to each of these mutants, P_*sigP*_-*lacZ* expression was no longer constitutive but was induced in the presence of cefoxitin (Fig. S8). This indicates that the *rsiP* mutations were responsible for the increased P_*sigP*_-*lacZ* expression.

**Figure 5.**
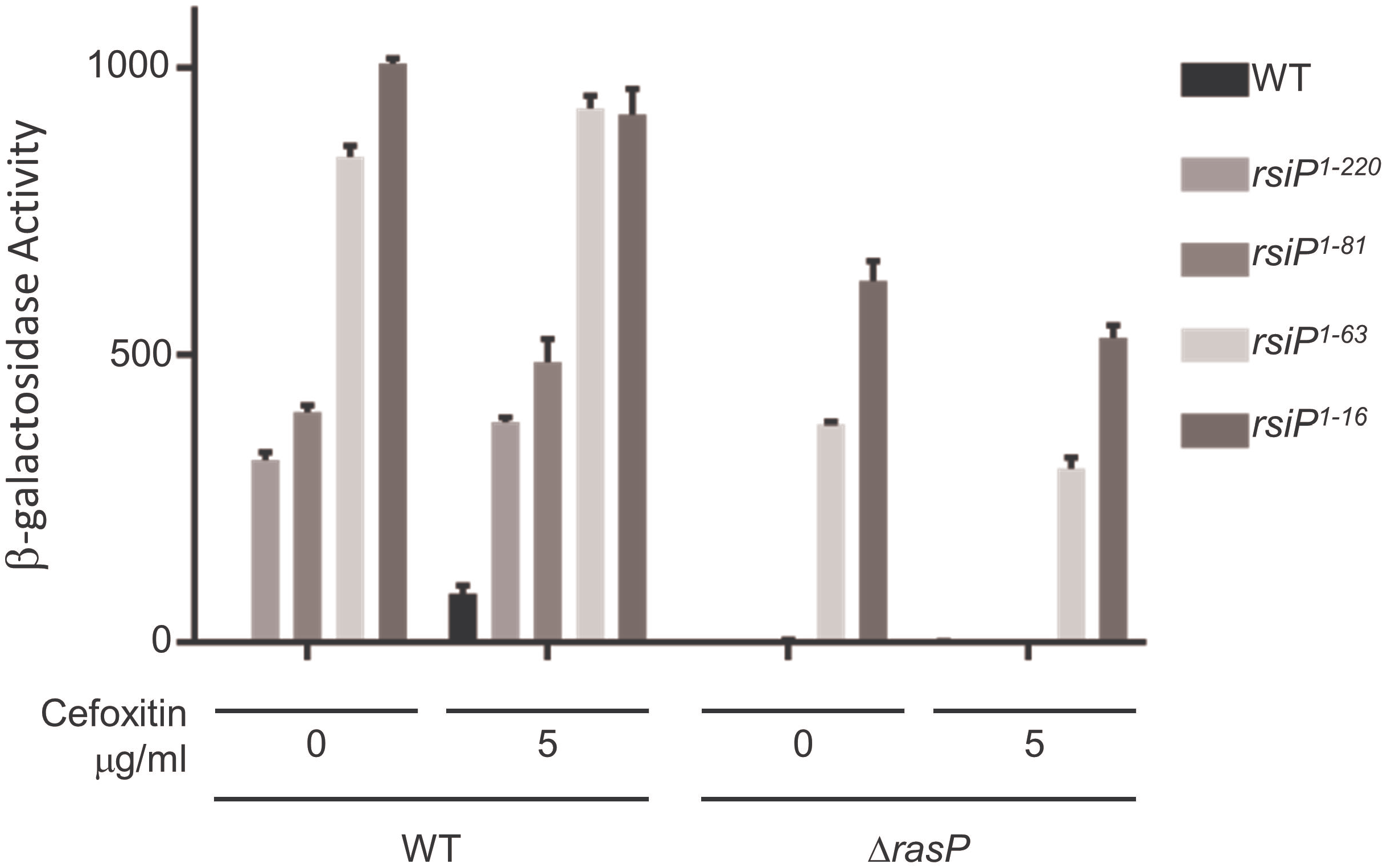
Truncations of RsiP lead to constitutive σ^P^ activation. To determine if RasP was required for ampicillin-inducible P_*sigP*_-*lacZ* expression we assayed β-galactosidase activity of *B. thuringiensis* with a transcriptional fusion P_*sigP*_-*lacZ* and different *rsiP* truncation mutants (WT, THE2549; RsiP^1-220^, THE2602; RsiP^1-80^, THE2628; RsiP^1-61^, THE2637; RsiP^1-16^, THE2642) and a Δ*rasP* deletion (WT, EBT140; RsiP^1-220^, EBT116; RsiP^1-80^, EBT148; RsiP^1-61^, EBT133; RsiP^1-16^, THE2605). Cells were grown overnight at 30°C and subcultured in LB and grown to OD_600_ of ~0.8 before being incubated with cefoxitin (5 μg/ml) for 1 hour. The experiment was performed in triplicate and standard deviation is represented by error bars.

In the σ^V^ and σ^W^ systems, RasP cleaves the anti-σ factors RsiW and RsiV within the transmembrane domain to activate the cognate σ factors (15, 35). The RsiP transmembrane is predicted to be residues 54-71 based on TMHMM (44). Two of the four RsiP truncations produce proteins with the transmembrane domain intact while the remaining RsiP truncations lack the transmembrane domain. Since RasP is known to cleave proteins within the transmembrane domain we hypothesized that those truncations which still contain a transmembrane domain would require RasP in order to activate σ^P^. To test this, we introduced the *ΔrasP* mutation into each of the *rsiP* mutants. In the absence of RasP, strains containing truncations which have a transmembrane domain (RsiP^1-220^, RsiP^1-80^; Fig. 4 and Fig. S7) no longer constitutively activate σ^P^ (Fig. 5). However, the strains with the *rsiP* truncation lacking the transmembrane domain (RsiP^1-16^, RsiP^1-61^) constitutively activate σ^P^ even in the absence of RasP (RsiP^1-16^, RsiP^1-61^; Fig. 4 and S5). Thus, RasP is required for σ^P^ activation when the transmembrane domain of RsiP is intact, consistent with the role of RasP as a site-2 protease.

### RasP cleaves within the transmembrane domain of RsiP and is not the regulated step in σ^P^ activation

In the case of σ^W^ and σ^V^ the rate-limiting step in σ factor activation is site-1 cleavage (15, 35). Since the identity of the site-1 protease is not currently known we sought to determine if RasP cleavage of RsiP is a rate-limiting step in σ^P^ activation. To test this, we constructed truncations of GFP-RsiP that lack the extracellular portion of RsiP. One truncation includes the transmembrane domain (*gfp-rsiP^1-72^*) and one truncation lacks the transmembrane domain (*gfp-rsiP^1-53^*). We expressed the truncated GFP-RsiP proteins in wild type and Δ*rasP* backgrounds and exposed these strains to cefoxitin (5 μg/ml). In wild type strains we found both GFP-RsiP^1-72^ and GFP-RsiP^1-53^ were degraded (Fig. 6 and Fig. S9). However, in the Δ*rasP* mutant the GFP-RsiP^1-72^ accumulated, while the GFP-RsiP^1-53^ was degraded (Fig. 6 and Fig. S9). These data indicate that GFP-RsiP^1-72^ requires RasP for degradation while GFP-RsiP^1-53^ does not. One possible interpretation is that GFP-RsiP^1-72^ is not produced or localized properly to the membrane. Thus, we confirmed that GFP-RsiP^1-72^ localizes to the membrane by fluorescent microscopy (Fig. S2). This suggests the RasP cleavage site of RsiP occurs within the transmembrane domain between amino acids 53 and 72. The presence or absence of cefoxitin had no effect on the degradation (Fig. 6 and Fig. S9). Since GFP-RsiP^1-72^ is constitutively degraded we conclude GFP-RsiP^1-72^ mimics the site-1 cleavage product and that RasP activity is not induced by cefoxitin. This suggests that RasP cleavage of RsiP is not the regulated step in σ^P^ activation and that site-1 cleavage is the step that is controlled by the presence of β-lactams.

**Figure 6.**
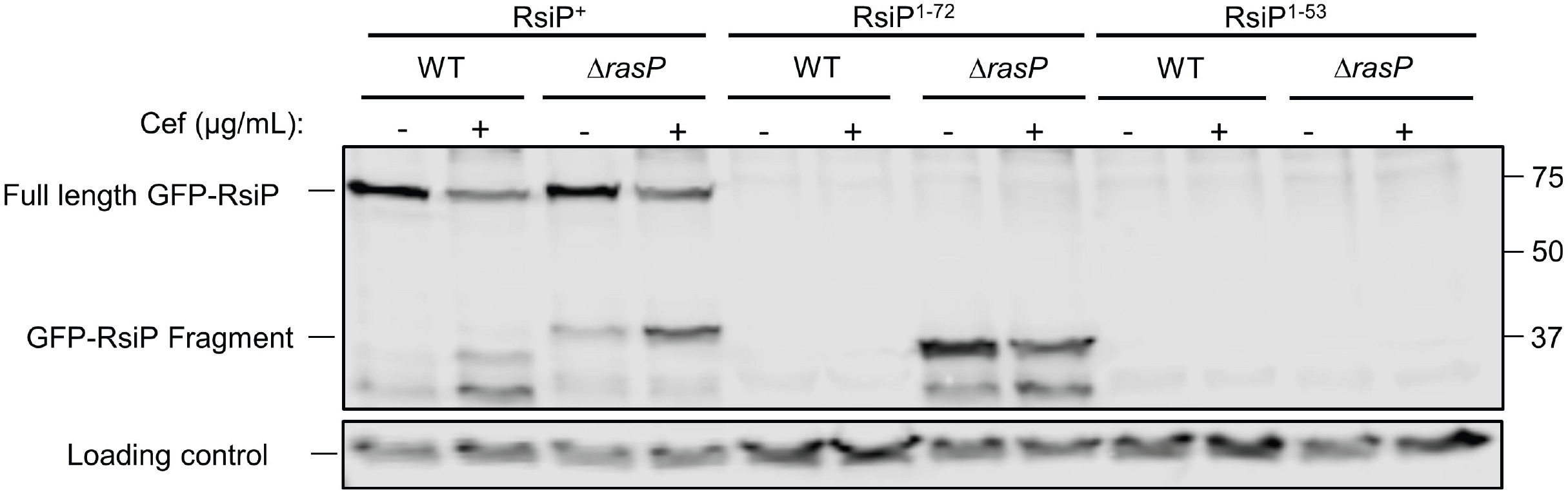
Truncation of RsiP results in constitutive degradation in a RasP-dependent manner. *B. thuringiensis* containing a tetracycline-inducible copy of *gfp-rsiP, gfp-rsiP^1-72^* (*rsiP* without the extracellular domain), or *gfp-rsiP^1-53^* (*rsiP* without the transmembrane and extracellular domains) were constructed in either wild type (*rasP*^+^) or a *ΔrasP* mutant strain [(GFP-RsiP-wild type (*rasP*^+^); EBT360), (GFP-RsiP *ΔrasP*; EBT366), (GFP-RsiP^1-53^ wild type (*rasP*^+^); EBT518), (GFP-RsiP^1-53^ *ΔrasP*; EBT510), (GFP-RsiP^1-72^ wild type (*rasP*^+^); EBT516), (GFP-RsiP^1-72^ *ΔrasP*; EBT533)]. Strains were subcultured 1:50 into LB supplemented with ATc (100 ng/ml) and grown to mid-log then incubated for 2 hours untreated (−) or treated (+) with cefoxitin (5 μg/ml) at 37°C. The immunoblot was probed with anti-GFP antisera. Streptavidin IR680LT was used to detect HD73_4231 (PycA homolog) which served as a loading control (63, 64). The color blot showing both anti-GFP and Streptavidin on a single gel is Fig. S9.

## Discussion

Many ECF σ factors are induced in response to extracytoplasmic stressors and initiate transcription of a subset of genes to modulate the cell’s response to these stresses. ECF σ factors can respond to signals such as misfolded periplasmic protein, antimicrobial peptides, or lysozyme. The ECF σ factors encoded in highly related organisms can vary widely. For example, *B. subtilis* encodes 7 ECF σ factors, while *B. thuringiensis* encodes 15 predicted ECF σ factors. The only ECF σ factor these organisms share in common is σ^M^ (45). Thus, there is a variability in how bacteria utilize ECF σ factors to respond to stress. Ross *et. al* demonstrated the novel ECF σ factor, σ^P^, is induced in the presence of ampicillin and initiates transcription of β-lactamases (5). Here we demonstrated σ^P^ responds specifically to a subset of β-lactams, while other β-lactams and cell wall targeting antibiotics fail to induce σ^P^ activation. We also showed σ^P^ confers varying degrees of resistance to these β-lactam antibiotics. We found that σ^P^ was not required for resistance to other cell wall antibiotics including vancomycin, nisin, bacitracin suggesting specificity in resistance to β-lactams and not a general cell envelope stress response.

For ECF σ factors to be activated, their cognate anti-σ factors must be inactivated. This can be accomplished via various mechanisms including: a conformational change of the anti-σ, partner switching where an anti-anti-σ factor frees the σ factor form the anti-σ factor, or proteolytic destruction of the anti-σ factor (Ho and Ellermeier, 2012; Helmann, 2016). The anti-σ factors RseA in *E. coli* as well as RsiW and RsiV in *B. subtilis* are degraded sequentially by regulated intramembrane proteolysis. Each of these anti-σ factors requires a different family of proteases to cleave the anti-σ factor at site-1 (14, 22, 30, 47, 48) while site-2 cleavage is carried out by the conserved site-2 protease (14, 15, 35). We hypothesize that σ^P^ is activated in a similar manner. Our data indicate σ^P^ is released from RsiP by proteolytic degradation when β-lactams are present. We found RasP is required for activation of σ^P^. We also observe that a RsiP degradation product approximately the size of our predicted RasP substrate accumulates in a Δ*rasP* mutant. This indicates RasP is required for degradation of RsiP. Our data also suggest, similar to other anti-σ factors, site-2 cleavage of RsiP is not the rate-limiting step since the C-terminal RsiP truncations are constitutively degraded and lead to constitutive σ^P^ activation in the absence of β-lactams. Thus, we hypothesize that RasP is required for site-2 cleavage of RsiP and an as yet unidentified protease is required to initiate degradation of RsiP by cleaving RsiP at site-1. We hypothesize, that like other ECF σ factors activated by regulated intramembrane proteolysis, site-1 cleavage of RsiP is likely the rate-limiting step in σ^P^ activation.

Our data suggest a subset of β-lactams induce σ^P^ activation. We found, in addition to ampicillin, σ^P^ is activated by cefoxitin, cefmetazole, cephalothin, cephalexin, and methicillin; but not by piperacillin, cefoperazone, cefsulodin, or antibiotics that target other steps in peptidoglycan biosynthesis. This raises the question: what is the signal for σ^P^ activation? The β-lactams could be sensed directly or indirectly. For example RsiV directly senses lysozyme and degradation of RsiV is rapid (31). In contrast activation of σ^E^ is indirect and due to buildup of products that occur when the outer membrane is damaged (31, 49). Our data suggest RsiP degradation is a relatively slow process. One possible interpretation of this is that β-lactam induced PG damage must accumulate to induce RsiP degradation. We hypothesize that the β-lactams we tested have different affinities for PBPs and this affinity may explain why some β-lactams induce σ^P^ while others do not. In other organisms, including *Streptococcus pneumoniae, B. subtilis* and *E. coli*, β-lactams can differentially target PBPs (50–52). This raises the possibility that activation of σ^P^ could be the result of inhibition of specific PBPs. Unfortunately, at this time we do not know which PBPs are targeted by the different β-lactams in *B. thuringiensis*. Thus, the precise mechanism and signal responsible for σ^P^ activation remain to be clearly defined.

## Materials and Methods

### Media and Growth Conditions

All *B. thuringiensis* strains are isogenic derivatives of AW43, a derivative of *B. thuringiensis* subspecies *kurstaki* strain HD73 (53). All strains and genotypes can be found in Table 3. All *B. thuringiensis* strains were grown in or on LB media at 30°C unless otherwise specified. Cultures of *B. thuringiensis* were grown with agitation in a roller drum. Strains containing episomal plasmids were grown in LB containing chloramphenicol (cam, 10 μg/ml) or erythromycin (erm, 10 μg/ml). *E. coli* strains were grown at 37°C using LB-ampicillin (amp, 100 μg/ml) or LB-cam (10 μg/ml) media. To screen for threonine auxotrophy, *B. thuringiensis* strains were patched on minimal media plates without or with threonine (50 μg/ml) (54, 55). The β-galactosidase chromogenic indicator 5-bromo-4-chloro-3-indolyl β-D-galactopyranoside (X-Gal) was used at a concentration of 100 μg/ml. Anhydrotetracycline (ATc, Sigma) was used at a concentration of 100 ng/ml.

**Table 3.**
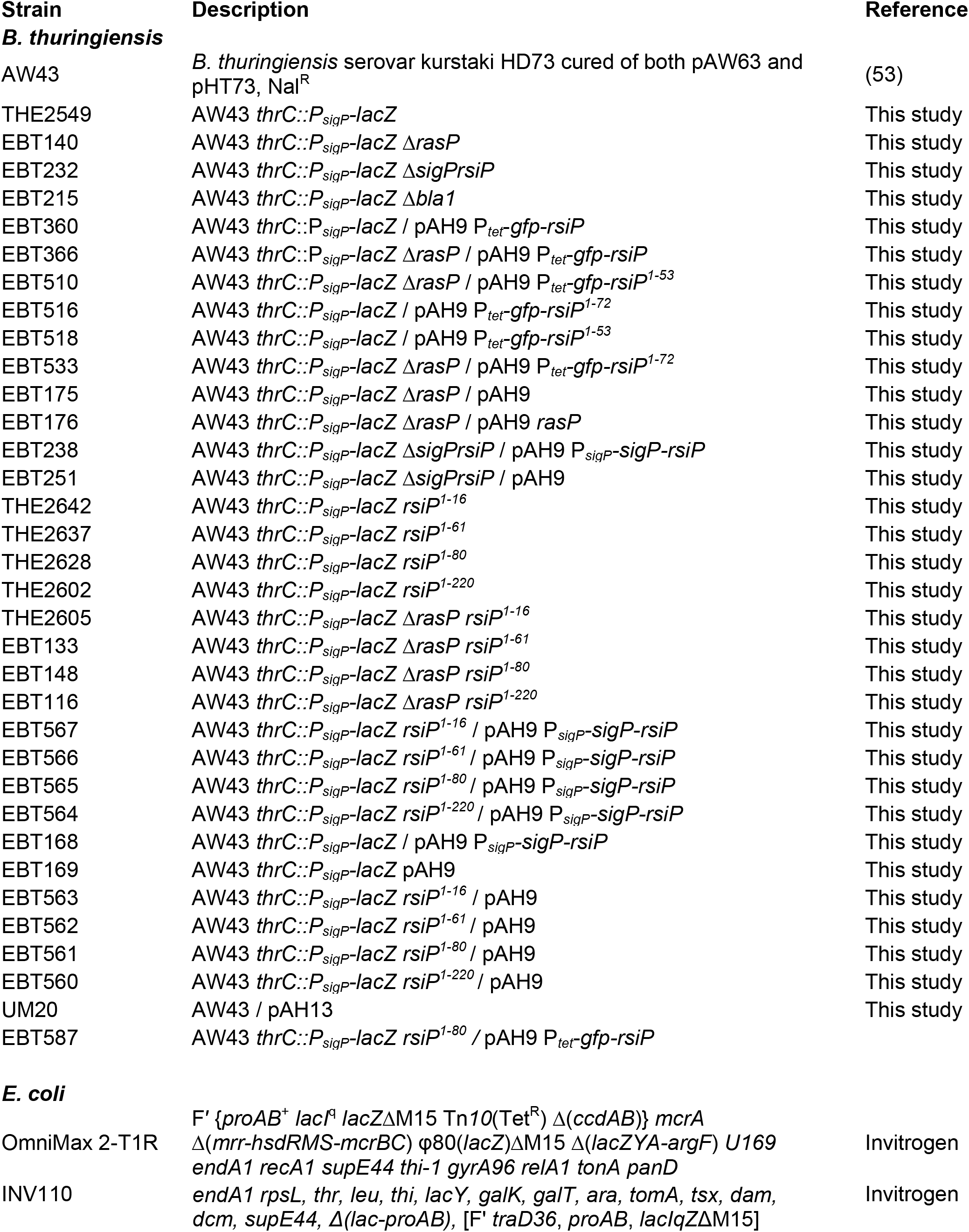
Strains.

### Strain and Plasmid Construction

All plasmids are listed in Table 4 which includes information relevant to plasmid assembly. Plasmids were constructed by isothermal assembly (56). Regions of plasmids constructed using PCR were verified by DNA sequencing. The oligonucleotide primers used in this work were synthesized by Integrated DNA Technologies (Coralville, IA) and are listed in Table S1. All plasmids were propagated using OmniMax 2-T1R as the cloning host and passaged through the non-methylating *E. coli* strain INV110 before being transformed into a *B. thuringiensis* recipient strain.

**Table 4.** Plasmids.

| Plasmid | Relevant features | Parent vector | Digest | Insert primers | Reference |
| --- | --- | --- | --- | --- | --- |
| pMAD | ori-pE194ts |  |  |  | Arnaud *et al* 2004 |
| pAH9 | ori-pE194 P<sub>sarA</sub>-*mcherry* |  |  |  | Malone *et al* 2009 |
| pAH13 | P<sub>tet</sub>-*gfp* |  |  |  | Malone *et al* 2009 |
| pRAN332 | P<sub>tet</sub>-*gfp* |  |  |  | Ransom *et al* 2015 |
| pEBT4 | ori-pE194ts, Δ*blaP* | pMAD | BglIII, EcoRI | 3832-3833; 3834-3835 | This study |
| pEBT5 | ori-pE194ts, Δ*rasP* | pMAD | BglIII, EcoRI | 3632-3633; 3634-3635 | This study |
| pEBT6 | ori-pE194ts, Δ*sigPrsiP* | pMAD | BglIII, EcoRI | 3776-3777; 3778-3779 | This study |
| pEBT13 | P<sub>tet</sub>-*gfp-rsiP* | pAH9 | HindIII, EcoRI | 3838-3839 | This study |
| pCE630 | P<sub>tet</sub>-*gfp-rsiP*<sup>1-72</sup> | pAH9 | HindIII, EcoRI | 3838-4258 | This study |
| pCE632 | P<sub>tet</sub>-*gfp-rsiP*<sup>1-53</sup> | pAH9 | HindIII, EcoRI | 3838-4259 | This study |
| pTHE960 | P<sub>sigP</sub>-*sigP*<sup>+</sup>*rsiP*<sup>+</sup> | pAH9 | HindIII, EcoRI | 3774-3775 | This study |
| pIA02 | P<sub>sarA</sub>-*rasP*<sup>+</sup> | pAH9 | EcoRI, KpnI | 3744-3745 | This study |
| pTHE946 | pE194ts | pMAD | BamHI, StuI |  | This study |
| pTHE948 | pE194ts, '*thrC thrB*' | pTHE946 | Scal, Sall | 2917-2918; 2919-2920 | This study |
| pTHE950 | pE194ts, '*thrC lacZ thrB*' | pTHE948 | XhoI, SbfI | 2922-2923 | This study |
| pTHE949 | pE194ts, '*thrC P<sub>sigP</sub>-lacZ thrB*' | pTHE950 | XhoI, Sall | 2929-2930 | This study |

To construct deletion mutants, we cloned 1 kb DNA upstream and 1 kb downstream of the site of desired deletion using primers listed in Table S1 onto the temperature sensitive pMAD plasmid (erythromycin-resistant) between the BglII and EcoRI sites (57).

Complementation constructs were constructed in pAH9 which is an *E. coli*-Gram positive shuttle vector with a pE194 origin of replication (58). Chromosomal DNA including the promoter sequence was cloned for *P_sigP_-sigP*^+^*rsiP*^+^ and cloned into pAH9 digested with EcoRI and HindIII while *rasP* was cloned downstream of the P_*sarA*_ promoter from *Staphylococcus aureus* by digesting with EcoRI and KpnI. In *B. thuringiensis* P_*sarA*_ has moderate constitutive expression.

To generate strains containing the *sigP* promoter fused to the *lacZ* reporter integrated into the chromosome, we constructed a number of intermediate vectors. To switch the antibiotic resistance of the temperature-sensitive pMAD vector, we constructed pTHE946 which contains the *E. coli* origin (ColE1 ori) of replication, erm-resistance gene (for selection in Gram-positives), amp-resistance gene (for selection in *E. coli* strains) and the temperature-sensitive origin (pE194 ori) from pMAD (7.3 kb StuI, BamHI fragment) as well as the conjugation origin of transfer and cam-resistance gene from pRPF185 (SmaI, BamHI fragment). The *thrC* (primers 2917, 2918) and *thrB* (primers 2919, 2920) genes were cloned into the ScaI, SalI digested pTHE946 plasmid (lacking *erm^R^* and *amp^R^* genes) to generate a vector (pTHE948) which can integrate into the *thrC* operon. A promoterless *lacZ* fragment (primers 2922, 2923) was added between the *thrC* and *thrB* genes of pTHE948 (XhoI, SbfI) to generate pTHE950. This plasmid (XhoI, NotI digested) was used to clone the *sigP* promoter (primers TE2929, 2930) to generate the P_*sigP*_-*lacZ* promoter fusion (pTHE949).

### *B. thuringiensis* DNA Transformation

Plasmids were introduced into *B. thuringiensis* by electroporation (59, 60). Briefly, recipient cells were grown to late-log phase at 37°C. For each transformation, cells (1.5 ml) were pelleted by centrifugation (9,000 x *g*) and washed twice in room temparature sterile water. After careful removal of all residual water, 100 μl of sterile 40% PEG 6000 (Sigma) was used to gently resuspend cells. Approximately 2-10 μl of unmethylated DNA (>50 ng/μl) was added to cells and transferred to a 0.4 cm gap electroporation cuvette (Bio-Rad). Cells were exposed to 2.5kV for 4-6 msec. LB was immediately added and cells were incubated at 30°C for 1-2 hours prior to plating on selective media.

### Construction of deletions or promoter-*lacZ* fusions in *B. thuringiensis*

To generate unmarked mutants and *thrC::P_sigP_-lacZ* strains, we used plasmid vectors containing the temperature-sensitive origin of replication (pE194 ori) from the pMAD plasmid (57). At permissive temperatures (30°C), pMAD replicates episomally as a plasmid. At non-permissive temperatures (42°C) pMAD must integrate into the chromosome via homologous recombination otherwise the plasmid will be lost to segregation and the strain will become sensitive to erythromycin. Plasmids were transformed into a *B. thuringiensis* recipient strain and selected for on LB-erm agar at 30°C. To select for the integration of the deletion plasmid into the recipient strain genome, plasmid-containing bacteria were grown at 42°C on LB-erm plates. The plasmid-integrated strain was then struck on LB agar at 30°C twice. Individual colonies were patched on LB and LB-erm agar to identify the erm-sensitive bacteria which had lost the deletion plasmid by segregation. To verify each deletion, genomic DNA was isolated from each strain candidate and PCR was used to verify the deletion. Integration of P_*sigP*_-*lacZ* fusion into the *thrC* operon results in threonine auxotrophy and can be identified by lack of growth on minimal media plates without threonine.

### Zones of Inhibition and Zones of Induction

To determine the zones of inhibition and induction by various antibiotics, we first washed mid-logarithmically grown cells in fresh LB. Washed cells were diluted 1:100 in molten LB agar containing X-gal (100 μg/ml) and poured into empty 100 mm Petri dishes. Sterile cellulose disks (8 mm) were saturated with different antibiotics and allowed to dry for greater than 10 minutes. After each antibiotic disk was placed on the solidified agar, plates were incubated at 30°C overnight before observing.

### β-Galactosidase Assays

To quantify expression from the *sigP* promoter, we measured the β-galactosidase activity of cells containing a P_*sigP*_-*lacZ* promoter fusion. Overnight cultures were diluted 1:50 in fresh LB media and incubated for 3 hours at 30°C. One ml of each subculture was pelleted (9,000 x *g*), washed (in LB broth) and resuspended in 1 ml LB broth lacking or including specified antibiotics. After 1 hour of incubation at 37°C, 1 ml of each sample was pelleted and resuspended in 200 μl of Z-buffer. Cells were permeabilized by mixing with 16 μl chloroform and 16 μl 2% sarkosyl (26, 61). Permeabilized cells (100 μl) were mixed with 10 mg/ml ortho-Nitrophenyl-β-Galactoside (ONPG, RPI, 50 μl) and OD_405_ was measured over time using an Infinite M200 Pro plate reader (Tecan). β-Galactosidase activity units (μmol of ONP formed min^−1^) X 10^3^/(OD_600_ X ml of cell suspension) were calculated as previously described (62). Experiments were performed in triplicate with the mean and standard deviation shown.

### Minimum Inhibitory Concentration Assay

To determine the minimum inhibitory concentration (MIC) for various antibiotics, we diluted overnight cultures of bacteria (washed in LB) 1:1000 in media containing two-fold dilutions of each antibiotic. All MIC experiments were performed in round-bottom 96-well plates. Each experiment was performed in triplicate and allowed to incubate for 24 hours at 37°C before observing growth or no growth.

### Immunoblot Analysis

Samples were electrophoresed on a 15% SDS polyacrylamide gel and proteins were then blotted onto a nitrocellulose membrane (GE Healthcare, Amersham). Nitrocellulose was blocked with 5% Bovine Serum Albumin (BSA) and proteins were detected with either 1:10,000 anti-GFP or 1:5,000 anti-RsiP^76-275^ anti-sera. Streptavidin IR680LT (1:10,000) was used to detect two biotin-containing proteins, PycA (HD73_4231) and AccB (HD73_4487), which serve as loading controls (63, 64). To detect primary antibodies, the blots were incubated with 1:10,000 Goat anti-Rabbit IR800CW (Li-Cor) and imaged on an Odyssey CLx Scanner (Li-Cor) or Azure Sapphire (Azure Biosystems). All immunoblots were performed a minimum of three times with a representative example shown.

## Acknowledgements

This work was supported by the Department of Microbiology and Immunology at the University of Iowa. We would like to thank Theresa Koehler for strains and advice. We also thank Leyla Slamti for protocols and members of the Ellermeier and Weiss labs for helpful comments.

---

**Supplemental Figure 1. GFP-RsiP is functional.** *B. thuringiensis rsiP^1-80^* (THE2628) containing tetracycline-inducible *gfp-rsiP* (EBT587) or empty vector (EBT561) were plated on LB Xgal without or with ATc (50 ng/ml).

**Supplemental Figure 2. GFP-RsiP localizes to the membrane.** *B. thuringiensis* expressing tetracycline-inducible *gfp-rsiP* (EBT360), tetracycline inducible *gfp-rsiP^1-72^* (EBT533), or empty vector (EBT169) were subcultured 1:50 with ATc 100 ng/mL and grown at 30°C to late log phase. 2 μL were spotted on a 1% agarose pad for immobilization and imaged for GFP localization. Phase-contrast and fluorescence micrographs were recorded on an Olympus BX60 microscope with a 100 UPlanApo objective (numerical aperture, 1.35). For the GFP micrographs a filter set from Chroma Technology Corp (catalog no. 41017) was used. The GFP filter consists of a 450-to 490-nm excitation filter, a 495-nm dichroic mirror (long pass), and a 500-to 550-nm emission filter. Micrographs were captured with a Hamamatsu Orca Flash 4.0 V2 complementary metal oxide semiconductor (CMOS) camera.

**Supplemental Figure 3. RsiP levels decrease in the presence of cefoxitin.** *B. thuringiensis* expressing tetracycline-inducible *gfp-rsiP* (EBT360) or empty vector (EBT169) was subcultured 1:50 into LB supplemented with ATc (50 ng/ml). At mid-log cells were incubated with (A) 5 μg/ml of cefoxitin for various times (0, 15, 30, 60, 120, or 180 minutes) or (B) increasing concentrations of cefoxitin (0, 0.05, 0.5, 5, 50, 500 μg/ml) for 1 hour. The immunoblot was probed with antisera against RsiP (α-RsiP^76-275^) followed by goat-anti-rabbit IgG IR800cw (Green). EV is wild type with pAH9 (EBT169) and GFP is wild type with pAH13 (UM20). Streptavidin IR680LT (Red) was used to detect HD73_4231 (PycA homolog) which served as a loading control (63, 64).

**Supplemental Figure 4. RsiP levels decrease in the presence of cefoxitin.** *B. thuringiensis* expressing tetracycline-inducible *gfp-rsiP* (EBT360), empty vector (EBT169) or GFP alone (UM20) were subcultured 1:50 into LB supplemented with ATc (50 ng/ml). At mid-log cells were incubated with increasing concentrations of cefoxitin (0, 0.05, 0.5, 5, 50, 500 μg/ml) for 1 hour. The immunoblot was probed with antisera against GFP (α-GFP) followed by goat-anti-rabbit IgG IR800cw (Green). Streptavidin IR680LT (Red) was used to detect HD73_4231 (PycA homolog) which served as a loading control (63, 64). The western is shown in (A) Black and white or (B) color.

**Supplemental Figure 5. Amino acid alignment of RasP.** An alignment of *B. subtilis* RasP and *B. thuringiensis* HD73_4301. The active site is marked by a red box.

**Supplemental Figure 6. RsiP degradation is dependent upon the site-2 protease, RasP.** *B. thuringiensis* containing a tetracycline-inducible copy of *gfp-rsiP*; wild type (EBT360) or *ΔrasP* (EBT366) were subcultured 1:50 into LB supplemented with ATc (50 ng/mL). At mid-log cultures were incubated for 1 hour untreated (−) or treated (+) with cefoxitin (5 μg/ml) at 37°C. The immunoblot was probed with antisera against GFP (anti-GFP) followed by goat-anti-rabbit IgG IR800cw (Green). EV is wild type with pAH9 (EBT169) and GFP is wild type with pAH13 (UM20). Streptavidin IR680LT (Red) was used to detect HD73_4231 (PycA homolog) which served as a loading control (63, 64).

**Supplemental Figure 7. RsiP mutants result in constitutive σ^P^ activity.** An alignment of RsiP mutants that result in nonsense, frameshift or point mutations. Amino acid residues in red are indicative of the change in amino acid sequence due to mutations. The mutation *rsiP^1-232^* was isolated 3 independent times while *rsiP^1-61^* and *rsiP^1-15^* were isolated twice each. The red box indicates the predicted transmembrane domain.

**Supplemental Figure 8. Complementation of RsiP mutants.** *B. thuringiensis* containing either Empty Vector (wild type, EBT169; *rsiP^1-220^*, EBT564; *rsiP^1-81^*, EBT565; *rsiP^1-61^*, EBT566; *rsiP^1-16^*, EBT567) or P_*sigP*_-*sigP-rsiP* (wild type, EBT168; *rsiP^1-220^*, EBT563; *rsiP^1-81^*, EBT562; *rsiP^1-61^*, EBT561; *rsiP^1-16^*, EBT560) were grown overnight and spotted onto LB X-gal (100 μg/ml) plates (A) lacking cefoxitin or (B) containing cefoxitin (5 μg/ml).

**Supplemental Figure 9. Truncation of RsiP results in constitutive degradation in a RasP-dependent manner.** *B. thuringiensis* containing a tetracycline-inducible copy of *gfp-rsiP, gfp-rsiP^1-72^* (*rsiP* without the extracellular domain), or *gfp-rsiP^1-53^* (*rsiP* without the transmembrane and extracellular domains) were constructed in either wild type of a *ΔrasP* mutant strain [(GFP-RsiP-wild type (*rasP*^+^); EBT360), (GFP-RsiP *ΔrasP*; EBT366), (GFP-RsiP^1-53^ wild type (*rasP*^+^); EBT518), (GFP-RsiP^1-53^ *ΔrasP*; EBT510), (GFP-RsiP^1-72^ wild type (*rasP*^+^); EBT516), (GFP-RsiP^1-72^ *ΔrasP*; EBT533)]. Strains were subcultured 1:50 into LB supplemented with ATc (100 ng/ml) and at mid-log were incubated for 2 hours untreated (−) or treated (+) with cefoxitin (5 μg/ml) at 37°C. The immunoblot was probed with antisera against GFP (anti-GFP) followed by goat-anti-rabbit IgG IR800cw (Green). Streptavidin IR680LT (Red) was used detect HD73_4231 (PycA homolog) which served as a loading control (63, 64).

**Supplemental Table 1. Oligonucleotides.**

